# Succinate Dehydrogenase (SDH)-subunit C regulates muscle oxygen consumption and fatigability in an animal model of pulmonary emphysema

**DOI:** 10.1101/2021.01.22.427763

**Authors:** Joseph Balnis, Lisa A. Drake, Catherine E. Vincent, Tanner C. Korponay, Diane V. Singer, David Lacomis, Chun Geun Lee, Jack A. Elias, Harold A. Singer, Ariel Jaitovich

**Author notes:** Correspondence and request for reprints should be addressed to Ariel Jaitovich MD, Albany Medical College, 47 New Scotland Avenue, Albany, NY. **Authors contributions:** JB, LAD, CEV, TCK, DVS, and AJ designed and performed experiments; JB and CEV performed proteomic analyses; LAD performed immunoblotting experiments; DL performed electron microscopy analysis; JB, DL, JAE, CGL, HAS and AJ designed the experiments and wrote the current manuscript.

## Abstract

Patients with pulmonary emphysema often develop locomotor muscle dysfunction, which is independently associated with disability and higher mortality in that population. Muscle dysfunction entails reduced force-generation capacity which partially depends on fibers’ oxidative potential, yet very little mechanistic research has focused on muscle respiration in pulmonary emphysema. Using a recently established animal model of pulmonary emphysema-driven skeletal muscle dysfunction, we found downregulation of succinate dehydrogenase (SDH) subunit C in association with lower oxygen consumption and fatigue-tolerance in locomotor muscles. Reduced SDH activity has been previously observed in muscles from patients with pulmonary emphysema and we found that SDHC is required to support respiration in cultured muscle cells. Moreover, *in-vivo* gain of SDH function in emphysema animals muscles resulted in better oxygen consumption rate (OCR) and fatigue tolerance. These changes correlated with a larger number of relatively more oxidative type 2-A and 2X fibers, and a reduced amount of 2B fibers. Our data suggests that SDHC is a key regulator of respiration and fatigability in pulmonary emphysema-driven skeletal muscles, which could be impactful to develop strategies aimed at attenuating this comorbidity.

## Introduction

Patients with chronic obstructive pulmonary disease (COPD) often develop locomotor muscle dysfunction, which entails atrophy and reduced force-generation capacity^1^. Muscle dysfunction, which occurs more often in patients with pulmonary emphysema than chronic bronchitis^2^, is strongly associated with higher mortality and other poor outcomes^3–6^. These associations persist even after adjusting for the level of pulmonary disease and other covariables suggesting that muscle dysfunction could be independently responsible for the worse prognosis^5,6^. Moreover, very few interventions can improve muscle status in pulmonary emphysema^7^, and none have shown mortality benefits^8^. It is generally accepted that maximal force-generation capacity predominantly depends on muscle mass, and submaximal force, or endurance, on fibers metabolic properties^1,9–11^. Inferential models suggest that fibers metabolic disturbances are associated with higher mortality in these patients, even after correcting for the magnitude of muscle atrophy^12^. While metabolic dysfunction directly impacts fibers contractility and fatigue-tolerance^11,13,14^, very little mechanistic research has been conducted on fibers respiration and their functional repercussions in the context of pulmonary emphysema.

We have recently described an animal model of pulmonary emphysema-driven skeletal muscle dysfunction that recapitulates many of the features observed in pulmonary emphysema patients, including dysfunctional fibers respiration and contractility^15–17^. In an unbiased analysis of that animal’s muscle proteome, we discovered a downregulation of succinate dehydrogenase-subunit C (SDHC), which is associated with lower oxygen consumption and decreased fatigue-tolerance. These associations are all regulated under chronic exercise, which improves SDH expression and activity, oxygen consumption and fatigue-tolerance^15^.

SDH is a tetrameric iron-sulfur flavoprotein of the tricarboxylic acid cycle (TCA)^18^. The correlation between SDH expression and fibers oxidative capacity was established long ago^19,20^, and indeed COPD patients’ skeletal muscles have been shown to have a reduced SDH activity^21^ associated with lower oxygen consumption^22^ even if the substrate availability (succinate) is controlled^23^. Moreover, TCA dysfunction caused by lower SDH activity could compromise O_2_ consumption and ATP generation^24^, leading to higher lactate production and fatigability^24–26^. Both lactate generation and fatigability are relatively increased in COPD patients muscles under endurance exercise in comparison with healthy individuals^27^. SDH is the only TCA cycle enzyme that also participates in the electron transport chain as complex II, and previous evidence demonstrates that dysfunction of SDH subunits located at the ubiquinol binding site, such as SDHC, cause increased reactive oxygen species (ROS) formation^28^, which occurs in COPD muscles^29,30^.

While is not known if SDH dysfunction causes the clinical manifestations of COPD myopathy, in this study we hypothesized that downregulation of SDHC directly contributes to reduced oxygen consumption and fatigue tolerance in COPD skeletal muscles and conducted both loss-and gain-of-function studies to mechanistically interrogate that interaction.

## Materials and methods

### Animals

Experiments were conducted using CC10-rtTA-IL-13 (IL13^TG^) doxycycline-inducible transgenic mice that develop chronic lung remodeling reminiscent of pulmonary emphysema upon induction^31,32^. CC10-rtTA-IL-13 heterozygote animals were bred to C57BL/6 mice to obtain IL13^TG^ *and* wild-type (IL13^WT^) littermate controls. Both IL13^TG^ (emphysema) and WT (non-emphysema, used as control littermates) mice were provided 0.5g/L doxycycline in their drinking water along with sucrose (0.5 mg/ml), starting at 5 weeks of age for a total dose of 17 weeks. Male and female mice were used for the studies. Food and water were accessible *ad libitum* and a 12-hour light/dark cycle was maintained. Sampling of skeletal muscle was performed directly after euthanasia by cervical dislocation. The time elapsed between animals’ euthanasia and muscle procurement and freezing never exceeded 3 minutes. All the procedures involving animals were approved by the Albany Medical College Institutional Animal Care and Use Committee (IACUC 07001) and animals were handled according to the National Institutes of Health guidelines; and all methods were performed in accordance with the relevant guidelines and regulations as stated by the Journal and public agencies.

### Muscle Histology

At 17 weeks post doxycycline initiation (22-23 weeks of age), freshly procured EDL muscles were placed on saline moistened gauze in a 60 mm culture dish on ice until freezing. A metal cup containing isopentane was cooled in liquid nitrogen until crystals formed of isopentane at the bottom of the cup. Muscles were transferred to pre-cooled Tissue-Tek embedding cassettes (EMS, Hatfield, PA, 62520) which were dropped into the cooled isopentane, submerging the muscle for 1 minute. Muscle samples were then drained and dried on gauze pads at −20°C to remove all isopentane. Frozen muscles were adhered to the sample stage using a small amount of Tissue-Tek optimal cutting temperature (OCT) compound (EMS, Hatfield, PA, 62550) and sections were completed using a Leica CM1860 Cryostat (Wetzlar, Germany); 10 μm sections were obtained for further analysis.

### Muscle Fiber Typing Immunofluorescence

Muscle sections were fixed for 15 minutes in acetone at −20°C, and then left at room temperature to dry for 30 minutes. Blocking was performed using Mouse-on-mouse blocking reagent (Vector Labs, Burlingame, California) for 1 hour at room temperature. Sections were then incubated for 45 minutes at 37°C with the primary antibodies indicated in the table at the end of the figures. Three washes were then performed with PBS. The following secondary antibodies from Jackson ImmunoResearch Laboratories were added all at 1:250 and incubated for 45 minutes at 37°C: anti-mouse IgG2b-DyLight 405, anti-mouse IgG1-Alexa 488, anti-mouse IgM-Alex594, and anti-rabbit IgG-Alexa 647. Three washes were then performed with PBS. Samples were mounted with Ibidi Mounting Medium (Martinsried, Germany). Images were captured the same day using confocal microscopy (Leica SPE).

### Electron microscopy

EDL muscle samples were placed in M. J. Karnovsky fixative immediately after procurement. Samples were dehydrated, embedded, cut on 60 nm-thick sections, mounted on corresponding grids and imaged by the University of Pittsburgh EM core facility.

### RNA Extraction, cDNA Synthesis, and Quantitative RT-PCR

RNA from tibialis anterior (TA) muscle was extracted using NucleoSpin RNA kit (Machery-Nagel, Düren, Germany). The reason TA was used is due to the higher yield of RNA per sample relative to other muscles. cDNA was synthesized using Quantitect Reverse Transcriptase Kit (Qiagen). Quantitative RT-PCR was performed using iTaq Universal SYBR Green Supermix (Bio-Rad) on a CFX96 Real-time PCR detection system (Bio-Rad). Each sample was run in triplicates, and relative expression levels of transcripts of interest were calculated using the comparative Ct (ΔΔCt) method with glyceraldehyde-3-phosphate dehydrogenase (GAPDH) as a housekeeping gene. Primers were purchased from Integrated DNA Technologies (IDT, IA), and a list of their sequences is presented in **Supplementary Table 2**.

### Modified Seahorse^®^ XFp Mitochondrial Stress Test

At 17 weeks post doxycycline initiation (22-23 weeks of age), mouse EDL muscles were isolated tendon to tendon and respirometry analyses were done as previously established^33^. In short, the muscle was bisected using a sterile scalpel and thin, even sections were placed in Matrigel precoated wells of an 8 well Seahorse^®^ plate. The first and last wells were reserved as coating-only controls, and samples were run in triplicates. Media added to each well contained XF base DMEM including 15mM glucose, 10mM sodium pyruvate, and adjusted to pH 7.4 using sodium hydroxide; and plates were equilibrated in a CO_2_-free incubator for 1 hour before running. Rotenone and antimycin A were used as respiratory chain inhibitors following manufacturers-suggested concentrations. Data was normalized to the total amount of protein contained in each well as determined by BCA assay. This method has the advantage of causing minimal disruption to the cytosolic environment that is intrinsic to the membrane permeabilization step used in conventional protocols^34,35^, does not entail potential mitochondrial respiration biasing die to isolation/differential centrifugation^36^, and thus is highly recommendable to specifically interrogate mitochondrial respiration in vivo^33^.

### Succinate dehydrogenase (SDH) Activity

At 17 weeks post doxycycline initiation (22-23 weeks of age), activity of SDH was determined using a succinate dehydrogenase assay kit (Sigma-Aldrich, MAK197) following the manufacturers protocol using the mouse tibialis anterior (TA) muscle lysate. Absorbance measurements were collected using a Cytation 5 plate reader (BioTek, Winooski, VT).

### Isolated muscle fatigability

At 17 weeks post doxycycline initiation (22-23 weeks of age), muscle fatigability was determined using an isolated muscle contractility platform. Immediately following euthanasia, the extensor digitorum longus (EDL) muscle was surgically isolated from the mouse carefully and a suture was tied around the tendon as each end of the muscle before cutting and removal. Special care was taken to assure no muscle stretching or damage was caused during the procurement process. Once removed, the isolated muscle was placed in ice-cold Ringer’s solution supplemented with 5.5mM glucose at pH 7.4 adjusted as needed with sodium hydroxide and bubbled with carbogen. The muscle was then suspended between the isometric force transducer (Harvard Apparatus, Holliston, MA) and the platinum-stimulating electrode tissue support (Radnoti, Covina, CA; 160152), lowered into the 25-mL tissue bath (Radnoti, Covina, CA; 166026) containing the same solution, also bubbled slowly with carbogen, but this time at room temperature; and allowed to equilibrate for an additional 15 min. Muscle tension was increased until baseline tension started to increase. A single 1-Hz, 40-V stimulus was delivered with a Grass S-88 electrical stimulator, and the peak contraction was recorded. After a 30-s rest, voltage was increased by 10 V and delivered again, recording peak contraction. This process was repeated until no additional increase in the peak contraction force was observed. The optimal length of the muscle was then determined by slowly increasing the muscle tension and delivering a single, maximal voltage stimulus, as previously determined above, while recording the peak force. After a 30-s rest, tension was slightly increased, and another stimulus was delivered while recording peak contraction. This process was repeated until maximal peak contraction force was achieved. Subsequent pre-fatigue stimuli were delivered at 1, 10, 20, 30, 50, 80, 100, and 120 Hz, while recording the peak force at each point and allowed for 1-min rest between each stimulus. Immediately after pre-fatigue process a train of additional stimuli were delivered by stimulating the muscle with 20 Hz, 500-ms duration, 1 train/s, for 5 min while recording constantly. Ftigue resistance was determined by comparing the initial and the final peak contraction measured during the fatigue program. All data collection and analysis were completed using the PowerLabs 4/20T (ADInstruments, Colorado Springs, CO) amplifier and LabChart 7 software.

### Succinate quantification

At 17 weeks post doxycycline initiation (22-23 weeks of age), amount of succinate was determined using a succinate colorimetric assay kit (Sigma-Aldrich, MAK184) following the manufacturers protocol using the mouse tibialis anterior (TA) muscle lysate. Absorbance measurements were collected using a Cytation 5 plate reader (BioTek, Winooski, VT).

### Cells and siRNA transfection protocol

C2C12 cells were cultured in normal conditions with 5% fetal bovine serum (FBS) growth medium. Cells were cultured in 6-well plates or seahorse respirometry plates as experimentally required. Cells in each setting were transfected with either 10nM scramble (control) or 10nM mouse SDHC siRNA-SMARTpool (Dharmacon, L048315-01-0005) using Lipofectamine 3000 (Life Technologies, L3000) and following the manufacturers recommended protocols. After 48 hours the same process was repeated before analyzing or collecting cells 48 hours after that.

### In-vivo electroporation

At 9 weeks post doxycycline initiation (14-15 weeks of age) the hair on the back legs of both animals was removed with Nair while under anesthesia. While still anesthetized both of the animals’ TA/EDL muscles was directly injected with 40uL of hyaluronidase (0.3U/uL in 0.9% NaCl) in 10uL aliquots across the length of the muscle then allowed to incubate without anesthesia for 2 hours. After 2 hours the animal was anesthetized and 40uL of plasmid with or without SDHC (1ug/uL in 0.9% NaCl) was injected in 10uL aliquots across the length of the muscle. The right leg was injected with the SDHC-containing plasmid and the left leg of the same animal was injected with the empty vector (plasmid with no SDHC). Using an ECM 830 square-wave electroporation system (BTX, Holliston MA) and Tweezertrode platinum electrodes (BTX, Holliston MA) five 25ms pluses of 150V was delivered through the clamped leg of the mouse. After the electroporation of both sides was completed the mouse continued receiving doxycycline for 8 additional weeks before collection at 17 weeks post initiation of doxycycline (22-23 weeks of age).

### Western blotting

Muscles were lysed in lysis buffer^50^ using a bead beating homogenizer (VWR, 10158-558) with 100x protease and phosphatase inhibitor cocktail (ThermoFisher, 78429). Cells were lysed using cell lysis buffer (Cell Signaling, 9803S). All samples were centrifuged at 22000xg for 10 minutes at 4C and transferred to fresh tubes to remove insoluble debris. Protein concentration were determined with a BCA assay (ThermoFisher, 23227) and normalized prior to gel electrophoresis. Precast 10% or 4-20% mini-PROTEAN TGX 10-well gels (Bio-Rad, 4561034) were used for protein separation electrophoresis. Proteins were transferred to nitrocellulose membranes using a wet-transfer system (Bio-Rad, 1703930) and blocked in 5% dry milk for 2 hours. Immunoblotting was performed in 1% dry milk overnight before 1-hour incubation with secondary antibody and visualization with chemiluminescence. Antibodies and specific concentrations used can be found in the supplement.

### Statistics

Data are expressed as the mean +/− S.E.M. When results were compared to a reference value, we used a single sample *t* test; when comparisons were performed between two groups, significance was evaluated by a Student’s *t* test, and when more than two groups were compared, analysis of variance was used followed by the Dunnett test using GraphPad Prism software. Results were considered significant when *p* < 0.05. For the proteomic analysis, data was processed using MaxQuant software (version 1.6.2.3). Searches were performed against a target-decoy database (*Uniprot (mouse)*, www.uniprot.org, October 28, 2018). Searches were conducted using a 20 ppm precursor mass tolerance and a 0.04 Da product mass tolerance. A maximum of 2 missed tryptic cleavages were allowed. The fixed modifications specified were carbamidomethylation of cysteine residues. The variable modifications specified were oxidation of methionine and acetylation of the N-terminus. Within MaxQuant, peptides were filtered to a 1% unique peptide FDR. Characterized proteins were grouped based on the rules of parsimony and filtered to a 1% FDR. Label-free quantification was performed within MaxQuant using MaxLFQ. Missing values were imputed using the Perseus tool available with MaxQuant. Quantitative. For the proteomic analysis, quantitative data from each experiment was log_2_ transformed and mean-normalized across all tissues for each given protein. Significantly changing proteins were identified using a two-sided Student’s T-test in Excel.

## Results

### Inducible pulmonary emphysema leads to reduced respiration in skeletal muscle

We have previously reported that *IL13^TG^* (emphysema) mice develop reduced body and muscle weight, decreased muscle fibers cross-sectional area and force-generation capacity in comparison with *IL13^WT^* (wild type) mice^15,16^. We also observed a significant reduction of fatigue-tolerance of extensor digitorum longus (EDL) muscles from *IL13^TG^* mice compared with *IL13^WT^* counterparts as evaluated in the isolated contractility platform (**Figure 1-A**). As fatigue-tolerance is directly influenced by oxidative capacity^37^, we determined EDL muscle oxygen consumption rate (OCR), which was significantly reduced in *IL13^TG^* in comparison with *IL13^WT^* animals (**Figure 1-B**). Oxidative capacity is sometimes^37^, yet not always^13^, correlated with the abundance of type II over type I fibers in COPD. However, we found the number of different fiber types to be unaltered in our *IL13^TG^* animals in comparison with wild type counterparts (**Figure 1-C**). Our previous data indicates no significant difference in mitochondrial mass between *IL13^WT^* and wild type animals^15^. To investigate whether the observed respiratory changes were associated with conspicuous qualitative mitochondrial changes, we conducted a systematic electron microscopy analysis of four *IL13^TG^* animals and four wild type counterparts. *IL13^TG^* animals’ muscles did not demonstrate altered mitochondrial structure as reflected by vacuolization, inclusion bodies, edema, intermembrane separation or any other obvious morphologic change (**Figure 1-D**). These data suggested that the reduced respiratory capacity of *IL13^TG^* animals’ muscles could be due to processes operating at sub-organellar level.

**Figure 1:**
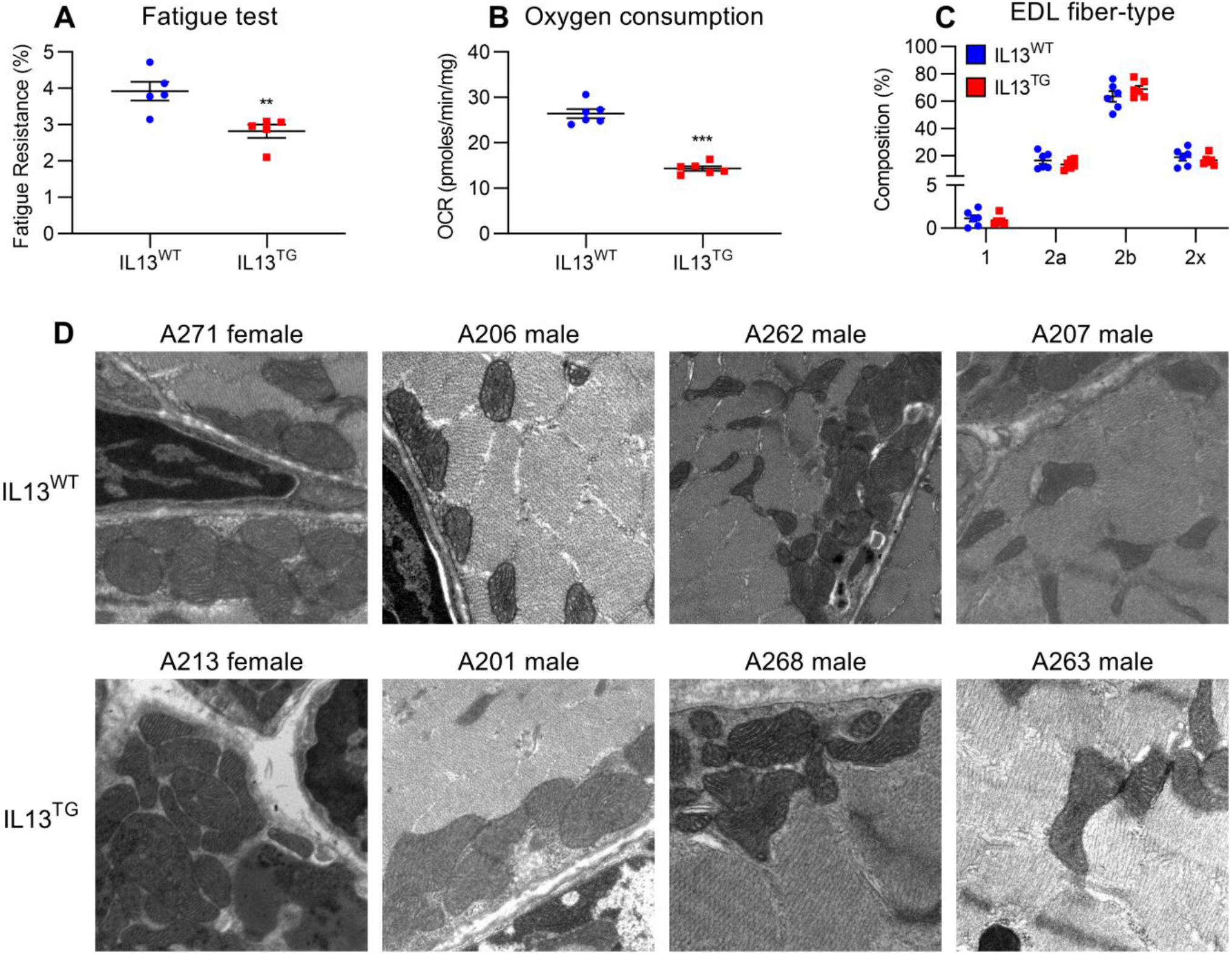
Inducible pulmonary emphysema leads to reduced respiration in skeletal muscle. Extensor digitorum longus (EDL) muscles from *IL13^TG^* (emphysema) and *IL13^WT^* (wild type) mice were ****A,**** assayed in the isolated contractility platform that demonstrated a reduced fatigue tolerance in *IL13^TG^* animals’ muscles (n=5). ****B,**** assayed in the plate respirometry (Seahorse^®^) platform, which demonstrated a reduced oxygen consumption rate (OCR) in *IL13^TG^* animals’ muscles (n=6). ****C,**** immuno stained with antibodies to detect myosin heavy chain (MyHC) isoforms in order to characterize fiber types, showing similar fiber type distribution in *IL13^TG^* and wild type mice (n=6). ****D,**** processed for electron microscopy evaluation, which ruled out any conspicuous structural mitochondrial abnormality (n=4; data above each panel designate the sex and number of the animal producing the micrography). *p*\*\*<0.01; ***<0.001.

### Inducible pulmonary emphysema leads to reduced expression of multiple proteins related to cellular respiration in skeletal muscle

To investigate potential mechanism leading to the respiratory phenotype of *IL13^TG^* animals’ muscles, we analyzed a previously published proteomic analysis of EDL muscles^15^. Ontology analysis of that dataset revealed a very significant downregulation of the bioenergetics-rich “ATP binding” term (see complete output dataset in **supplement** and reference^15^). We identified multiple dysregulated proteins that could account for the reduced oxygen consumption rate observed in *IL13^TG^* mice (**Figure 2-A**). Among them, succinate dehydrogenase subunit C was found particularly attractive given the previous description of its reduced activity in COPD patients muscles21 (**Figure 2-B**). Messenger RNA level of SDHC, but not of the other three subunits, was found downregulated in *IL13^TG^* animals’ muscles (**Figure 2-C**). Moreover, muscle SDH enzymatic activity assay demonstrated a significant reduction in *IL13^TG^* in comparison with wild type mice (**Figure 2-D**). As the enzyme activity can be limited by reduced substrate availability, we determined succinate concentration, which was increased in *IL13^TG^* versus wild type mice (**Figure 2-E**), suggesting that the reduced activity was not substrate-dependent.

**Figure 2:**
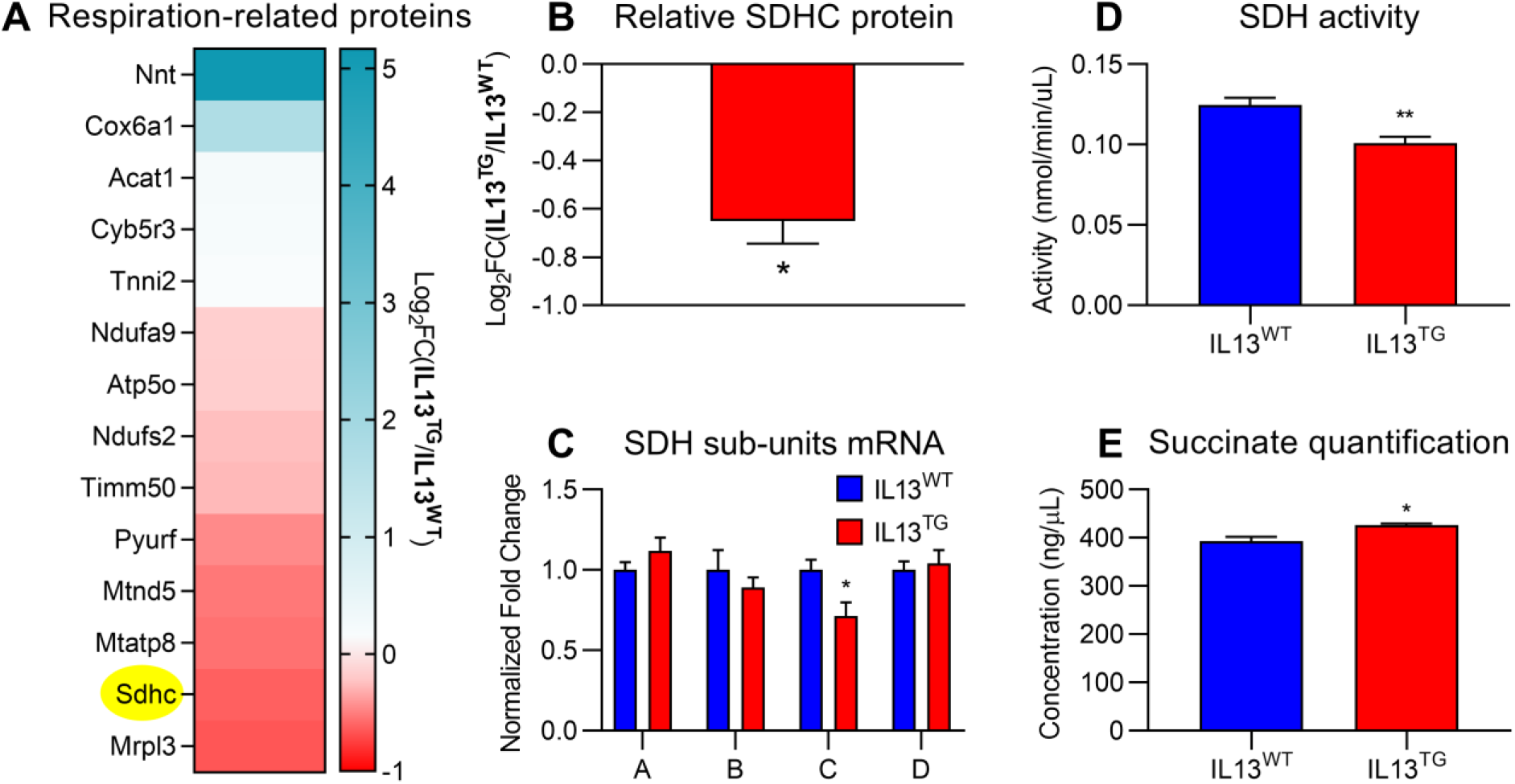
Inducible pulmonary emphysema leads to skeletal muscle reduced expression of multiple proteins related to cellular respiration. *A,* Proteomic analysis of EDL muscles identified potentially relevant associations of mitochondrial dysfunction in *IL13^TG^* versus wild type mice, including succinate dehydrogenase (SDH)-subunit C, which is highlighted in yellow; all the described proteins are significantly regulated after correction for multiple comparisons/false discovery rate; color scale denotes robustness of positive (blue) or negative (red) regulation; see full dataset at^15^. ****B,**** quantification of SDHC downregulation in *IL13^TG^* versus wild type mice captured in the proteomic analysis. ****C,**** RNA expression of SDH isoforms show that SDHC is the only component of the tetramer SDH/complex II that is significantly downregulated in COPD muscles. ****D,**** SDH activity is reduced in tibialis anterior (TA) muscle from *IL13^TG^* versus wild type mice. ****E,**** succinate is accumulated in TA muscles from *IL13^TG^* versus wild type mice, indicating that SDH activity *is not limited by substrate availability*. N=5; * *p*<0.05; \*\**p*<0.01.

### Downregulation of SDHC expression leads to reduced enzymatic activity and oxygen consumption rate in cultured muscle cells

While SDH activity is required for cellular respiration, the observed reduction in *IL13^TG^* mice muscles is rather modest (~25%), making possible that the remaining SDH activity is enough to support oxidative metabolism. To determine if the observed magnitude of SDHC downregulation is sufficient to cause reduced oxygen consumption, we used cultured non-transformed immortalized skeletal muscle C2C12 cells^38^. Experiments were calibrated based on previous data on SDHC half-life^39^ to reach a downregulation level in the range of what *IL13^TG^* mice demonstrate (**Figure 2-B**). Transfection of these cells with SDHC-specific siRNA led to a significant reduction of that subunit mRNA product and not the other three (**Figure 3-A**), which was not associated with evidence of cellular toxicity or altered morphology (**Figure 3-B**). SiRNA also led to a decreased protein product (**Figure 3-C**), enzymatic activity (**Figure 3-D**), and oxygen consumption rate (**Figure 3-E**). These data supported further experiments to define if SDHC downregulation in *IL13^TG^* mice muscles contributes to their altered respiratory profile.

**Figure 3:**
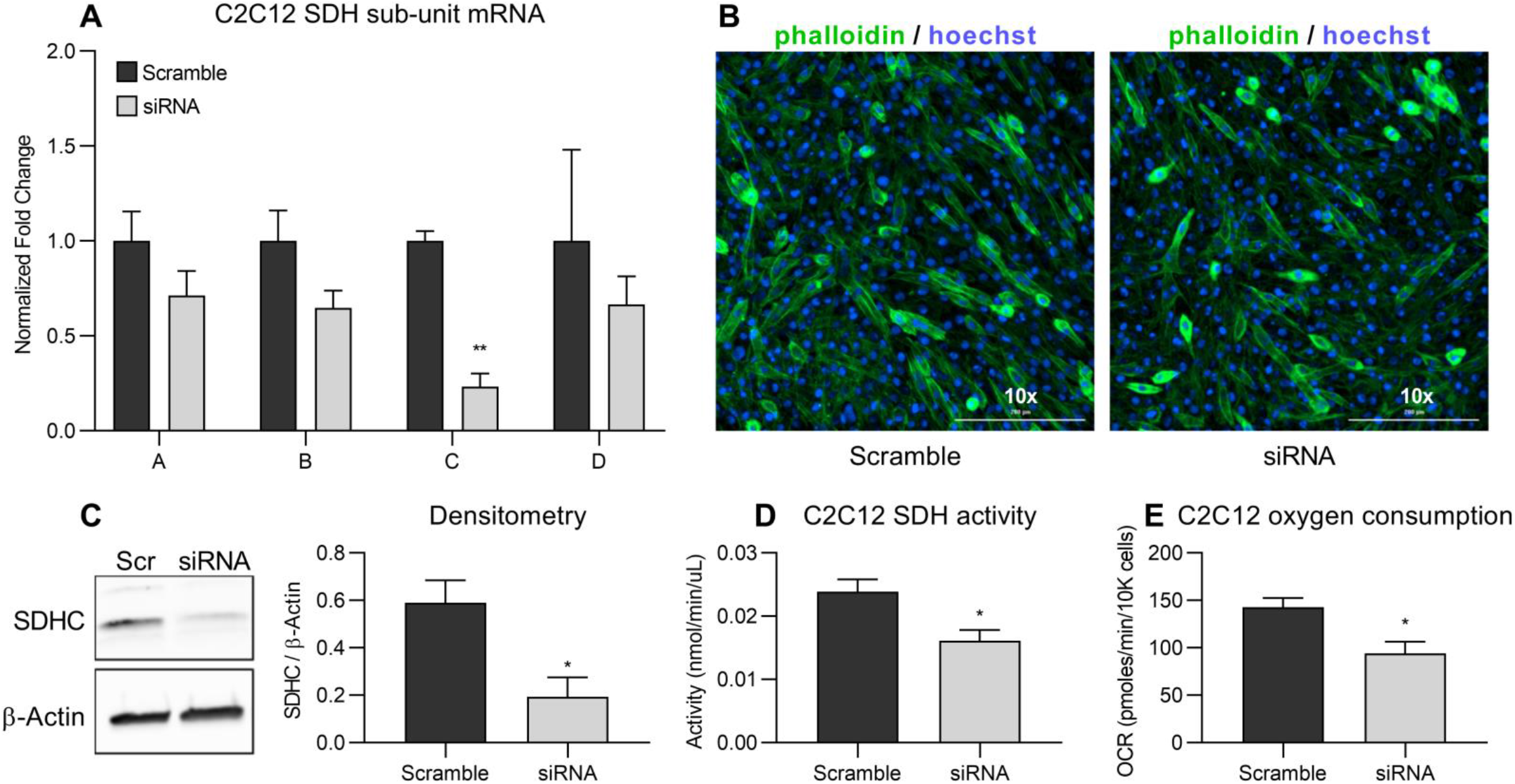
Downregulation of SDHC expression leads to reduced enzymatic activity and oxygen consumption rate. Cultured muscle (C2C12) cells were transfected with scramble and specific SDHC siRNA. A, qPCR demonstrated a significant reduction of SDHC mRNA, while the other subunits remained not significantly reduced. ****B,**** microscopic evaluation of cells stained with phalloidin and Hoechst indicates that SDHC downregulation does not lead to evidence of cellular toxicity. ****C,**** Western blotting using SDHC-specific antibody demonstrates downregulation of the protein product in siRNA specific-transfected cells. ****D,**** SDH enzymatic activity is reduced about 35-40% in cells transfected with specific SDHC siRNA; ****E,**** Seahorse^®^ platform-determined oxygen consumption rate (OCR) is reduced in cells previously transfected with specific SDHC siRNA. N=3; * *p*<0.05; \*\**p*<0.01.

### *In vivo* overexpression of SDHC in *IL13^TG^* mice skeletal muscles leads to improved respiration and fatigue-tolerance

We reasoned that if SDHC downregulation contributes to reduced muscle oxygen consumption and fatigue-tolerance, then SDHC restitution should attenuate these deficits. We then performed *in-vivo* muscle electroporation using SDHC-holding or empty (control) constructs. As electroporation causes local muscle injury and the complete myogenic response occurs in about one month^40^, we reasoned that performing experiments at two months post-electroporation would minimize the possible confounding effect of inadequate muscle repair. Moreover, post-injury muscle repair is known to cause upregulation of slower-twitch, and thus more oxidative phenotype^37,41,42^ which could potentially skew the baseline results. Thus, we performed both gain-of-function and control experiments on the same experimental animals, using their right and left TA muscles, respectively. We determined electroporation efficiency by green-fluorescent protein (GFP) reporting (**Figure 4-A-1**), which was found on both TA muscles; and SDHC sequence overexpression by FLAG immunoblotting, which only happened in the experimental (right) leg (**Figure 4-A-2**, also see construct map in **supplementary figure**). SDHC mRNA overexpression and SDH activity were also significantly elevated compared with empty vector-transfected contralateral muscles (**Figure 4-B** and **C**). Importantly, the succinate accumulation seen in *IL13^TG^* mice was not observed post-electroporation compared with empty vector-transfected contralateral TA muscle, also indicating a better SDH activity (**Figure 4-E**). We then performed tibialis anterior (TA) muscle plate respirometry (Seahorse^®^) assay and found that OCR values were significantly increased in the SDHC transfected leg when compared with the contralateral muscle (**Figure 4-D**). Moreover, to evaluate the functional effect of improved respiratory capacity, we determined EDL muscle fatigability on the isolated contractility platform, which showed that the reduced fatigue-tolerance of *IL13^TG^* mice was abrogated after SDHC overexpression (**Figure 4-F**). These data indicate that SDHC gain-of-function causes improved skeletal muscle oxygen consumption and fatigability in this animal model of pulmonary emphysema.

**Figure 4:**
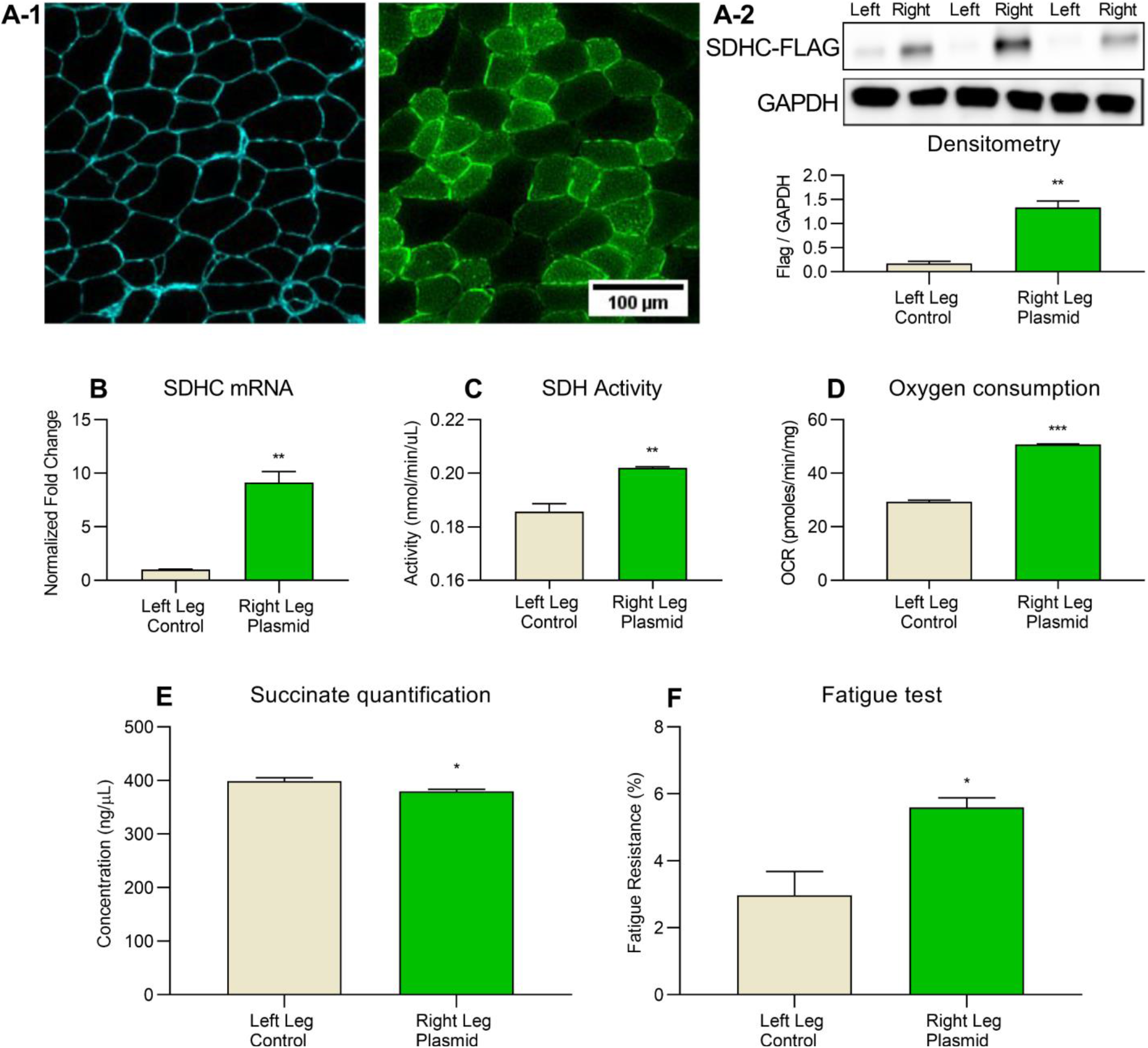
Overexpression of SDHC in *IL13^TG^* mice skeletal muscles leads to improved respiration and fatigue-tolerance. Muscles from *IL13^TG^* (emphysema) and *IL13^WT^* (wild type) mice were electroporated with a construct holding the SDHC-Flag sequence (right leg) and an SDHC-lacking vector (left leg). Both plasmids hold a green-fluorescent protein sequence as electroporation efficiency reporter. ****A-1,**** microscopic evaluation of muscle section demonstrating preservation of histoarchitecture and positive fluorescence post-electroporation; ****A-2,**** Western blotting of muscle samples demonstrating expression of Flag sequence in right leg versus negative expression in left leg. GAPDH was used as a lane loading control. ****B,**** SDHC mRNA normalized to GAPDH mRNA was over expressed in the right versus left TA muscle post-electroporation. ****C****, SDH activity was significantly elevated in in the right versus left TA muscle post-electroporation; ****D,**** oxygen consumption rate (OCR) was significantly elevated in the right versus left EDL muscle post-electroporation. ****E,**** succinate quantification demonstrated not significantly different levels between the right versus left TA muscle post-electroporation. ****F,**** fatigue resistance was significantly elevated in the right versus left EDL muscle post-electroporation. N=4; * *p*<0.05; \*\**p*<0.01; ***<0.001.

### Effect of SDHC overexpression on fiber type identity in *IL13^TG^* mice skeletal muscles

Fiber type as defined by the myosin heavy chain (MyHC) isoform expression correlates with the oxidative capacity and resistance to fatigue^37^. These profiles in mice transition in a spectrum that goes from slower to faster-twitch, and from more to less oxidative capacity and fatigue resistance as follows: type 1↔2A↔2X↔2B^43,44^. As improved mitochondrial biogenesis and respiratory capacity has been shown to cause type 2-to-1 fiber switch^45^, we interrogated whether SDHC overexpression had an effect on *IL13^TG^* mice fibers profile. We found that EDL muscles with overexpressed SDHC-holding constructs had a significant increase in type 2-A and 2X fibers and a reduction of 2B fibers; and no effect on type 1 fibers (**Figure 5**). These data suggest that the myosin isoform expression, at least in the presented model, could operate downstream of the elevation of oxidative capacity driven by SDHC normalization.

**Figure 5:**
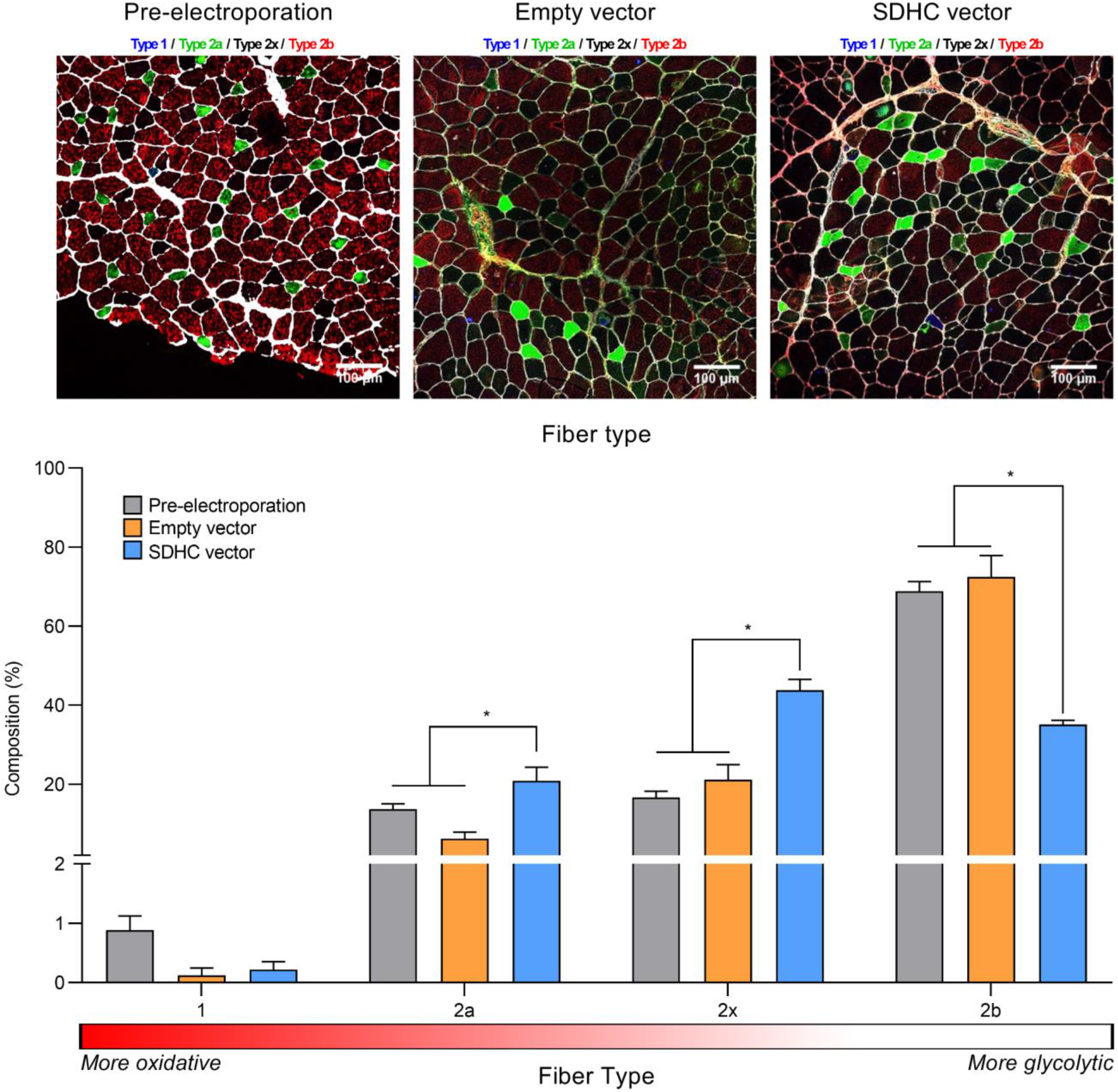
Effect of SDHC overexpression on fiber type identity in *IL13^TG^* mice skeletal muscles. Pre and post-electroporation EDL muscles from *IL13^TG^* mice were immuno stained with antibodies to detect myosin heavy chain (MyHC) isoforms in order to characterize fiber types. Post electroporation, there was an increase in relatively more oxidative type 2A and 2X fibers, and a reduction in relatively glycolytic type 2B fibers; without a significant change in the number of type I fibers, N=4; * *p*<0.05

## Discussion

While COPD locomotor muscles’ reduced endurance, fatigue-tolerance and oxidative capacity have been consistently observed^27,46^, there has been debate on whether intrinsic defects of mitochondrial respiration occur in that setting or not^23,30,47–49^. In this work, we used a recently described animal model of pulmonary emphysema-induced muscle dysfunction^15–17^ to interrogate the potential mechanisms underpinning abnormal fibers metabolism. The fact that pulmonary emphysema animal muscles demonstrate SDHC downregulation, along with previous observations indicating similar reduced SDH activity in humans^21,49,50^, led us to postulate that SDH dysfunction could be responsible for a reduced fibers respiration capacity. We confirmed a decreased SDH enzymatic activity in the TA muscles, which is associated with succinate accumulation, indicating that substrate availability is not rate-limiting the enzyme work. Moreover, siRNA driven SDHC silencing led to SDH loss of function in cultured muscle cells that was of similar magnitude to the one observed in emphysema mouse muscles. This led to a substantial decrease in oxygen consumption rate, supporting the hypothesis that SDHC downregulation *in-vivo* could be partially responsible for the metabolic and functional phenotype observed in emphysema mouse muscles. Indeed, in-vivo SDH gain-of-function in emphysema mouse muscles via SDHC overexpression led to improved oxygen consumption and fatigue-tolerance; and also, a reduction in succinate accumulation. This strongly suggests that SDHC downregulation is, at least partially, responsible for muscle metabolic dysfunction seen in this model of pulmonary emphysema. We believe that a chronic disease model such as *IL13^TG^*-induced muscle dysfunction is consistent with the modest, although statistically significant, reduction of SDH activity.

While most of the research conducted in the field of COPD-associated muscle dysfunction has so far focused on the conspicuous reduction of muscle mass^1,4–6,15–17,51–53^, to our knowledge this is the first mechanistic study that disaggregates the investigation of muscle metabolic properties in the context of pulmonary emphysema. While our study is focused on the fibers oxygen consumption rate and its effect on fatigue-tolerance, we speculate that other biologically relevant processes could also be involved and should be investigated in the future. Significantly, succinate accumulation could inhibit 2-oxoglutrate-dependent dioxygenases (2-OGDD)^54^, driving DNA and/or chromatin epigenetic changes including hypermethylation of regulatory elements^55–57^ creating a non-permissive state of critical genes needed to maintain muscle metabolic integrity. For example, it is known that in SDH-deficient cells, there is an increase in the fraction of methylated DNA (5-mC/5-hmC ratio) and chromatin which is associated with the silencing of key genes involved in differentiation^54,58^. Succinate accumulation could inhibit prolyl-hydroxylases (PHDs), which are 2-OGDD enzymes critical in the regulation of the transcription factor HIF-1 signaling, a master regulator of O_2_ homeostasis. Succinate has also been involved in post-translational protein modification (succinylation)^59^, and in cellular signaling via a specific receptor such as G protein-coupled receptor succinate receptor 1 (SUCNR1)^60^; all of which could amplify metabolic dysfunction and other processes. Interestingly, a recent study indicates that exogenous succinate supplementation to cultured muscle cells and mice undermines muscle regeneration after injury^61^.

While SDH catalyzes the tricarboxylic acid (TCA) cycle conversion of succinate to fumarate, it is also the only TCA enzyme that participates in the electron transport chain (ETC) as the complex II^62^, making it an attractive target to investigate the interaction between bioenergetics and oxidative stress in COPD muscles^63^. Importantly, previous evidence indicates that dysfunction of SDH subunits located at the ubiquinol binding site-such as subunit C-leads to increase in ROS formation whereas proximal ones such as SDHA do not^28^. Indeed, complex II-dependent increase of ROS has been demonstrated in COPD patients’ muscles despite succinate contribution^23^. Moreover, in an analysis of this COPD animal model muscle proteome we recently reported the upregulation of antioxidant ferroproteins, likely indicating ongoing activation of oxidative stress signaling^15,16^, and supporting the role of mitochondria-induced ROS generation in that setting. Future studies focused on the potential ROS-generation role of SDH dysfunction could also be investigated with gain-of-function analyses. The mechanism leading to SDHC downregulation in this animal model remains to be elucidated. While the *IL13^TG^* mice are hypoxemic^15–17^, we speculate that hypoxia is unlikely to be an upstream regulator of SDH-driven reduced respiration. Indeed, previous data on non-muscle cells^64^ and in rat skeletal muscle^65^ indicates that hypoxia does not cause significant downregulation of SDH^66^. Future studies could investigate the specific interaction between hypoxia and other signals with SDH in the regulation of muscle respiration and fatigability.

We found an increase in oxidative type 2A and 2X, and a reduction in the less oxidative 2B, in post SDHC overexpression EDL muscles from *IL13^TG^* mice. Although fiber type identity partially depends on the motoneuron innervation profile^37,67^, recent data indicates that it can also be regulated downstream of subcellular processes associated with mitochondrial respiration^45^. Moreover, it has been recently reported that myogenesis can contribute to fiber-specific profiles based on the recruitment of twist-2 positive myogenic progenitor cells^68^. Interestingly, we have shown that there are no significant baseline differences between fiber types in EDL muscle from *IL13^TG^* and *IL13^WT^* mice, yet overexpression of SDHC leads to larger number of types 2A and 2X fibers suggesting that muscle fiber type profile could be regulated by SDHC independently of its SDH complex enzymatic function. Indeed, previous evidence shows that SDHC has cellular functions that are independent of its canonical catalytic function^18^. Future research will be required to elucidate the mechanisms regulating that SDHC-induced fiber change in the context of *IL13^TG^* mice.

While to our knowledge this is the first study that mechanistically investigates muscle fiber oxygen consumption in pulmonary emphysema, we realize this study has limitations. First, the present data is based on an animal model that does not necessarily fully reflect the process taking place in patients. For instance, MyHC-2B is not detectable in humans^69^, which should be considered when extrapolating the present data to COPD myopathy. Validation of this observations in other animal models of COPD and human samples will be important to address this limitation in the future. Second, the SDH overexpression approach uses a standard in vivo electroporation technique. Although this is a validated method to generate loss- and gain of function settings, it cannot prevent biases associated with injury-repair and off target effects of larger-than-physiological protein expressions. Future studies involving a genetic animal model of inducible, muscle targeted SDHC over expression could better address this limitation.

## Conclusion

Succinate dehydrogenase subunit C (SDHC) overexpression abrogates reduced oxygen consumption and fatigue-tolerance observed in an animal model of pulmonary emphysema-induced skeletal muscle dysfunction, which is associated with an increase in relatively more oxidative type 2A and X fibers (**Figure 6**). This mechanism could stimulate research to support SDH function in COPD patients with skeletal muscle dysfunction^70^.

**Figure 6:**
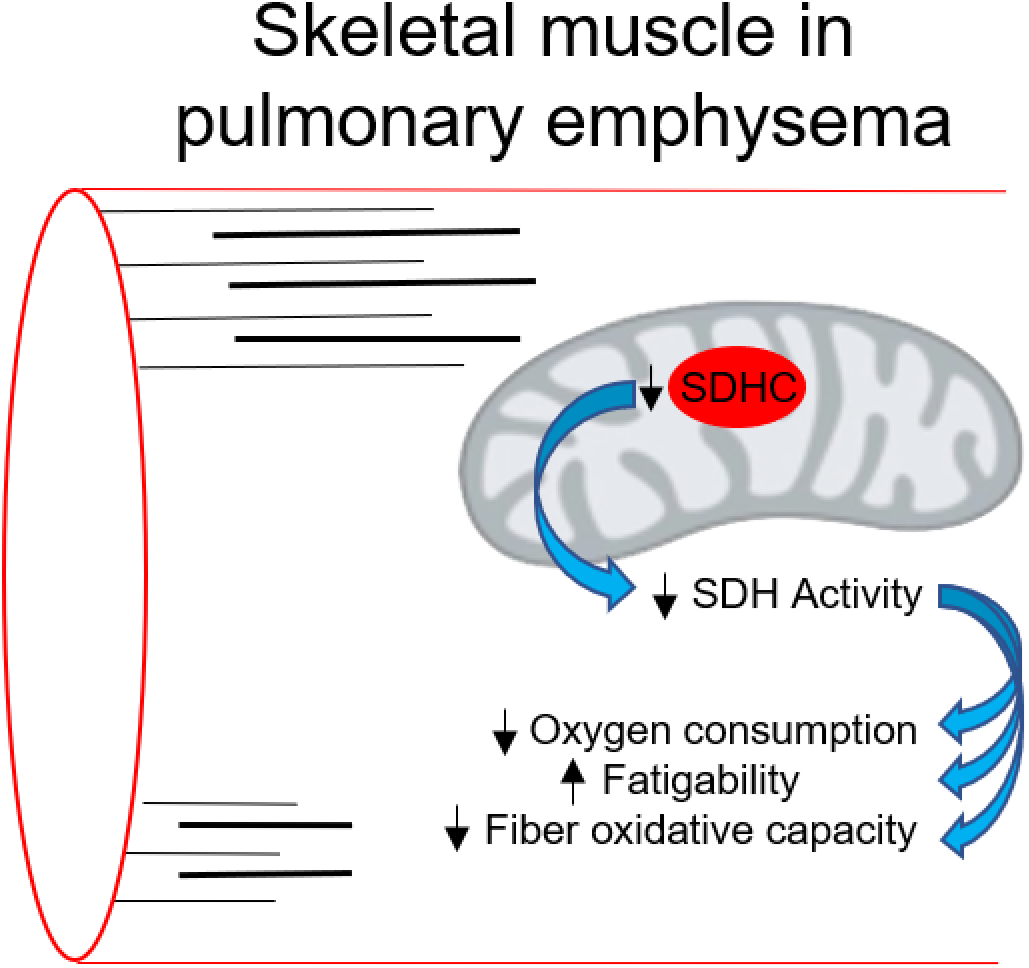
Cartoon illustrating the proposed mechanism leading to SDHC-driven reduced oxygen consumption rate and fatigue-tolerance in this model of pulmonary emphysema.

## Acknowledgements

We would like to acknowledge the National Center for Quantitative Biology of Complex Systems (P41 GM108538) at the University of Wisconsin, Madison, WI, for graciously providing the LC-MS/MS instrument time required to analyze our proteomic samples, and especially Dr. Joshua J. Coon for facilitating sample transport and data acquisition at the NCQBCS. We also thank Nina Martino and Alejandro P. Adam for their help generating the SDH-holding plasmid used in the in-vivo electroporation experiments.

---

**File S1: Proteomic analysis output file.**

**Figure S1:**
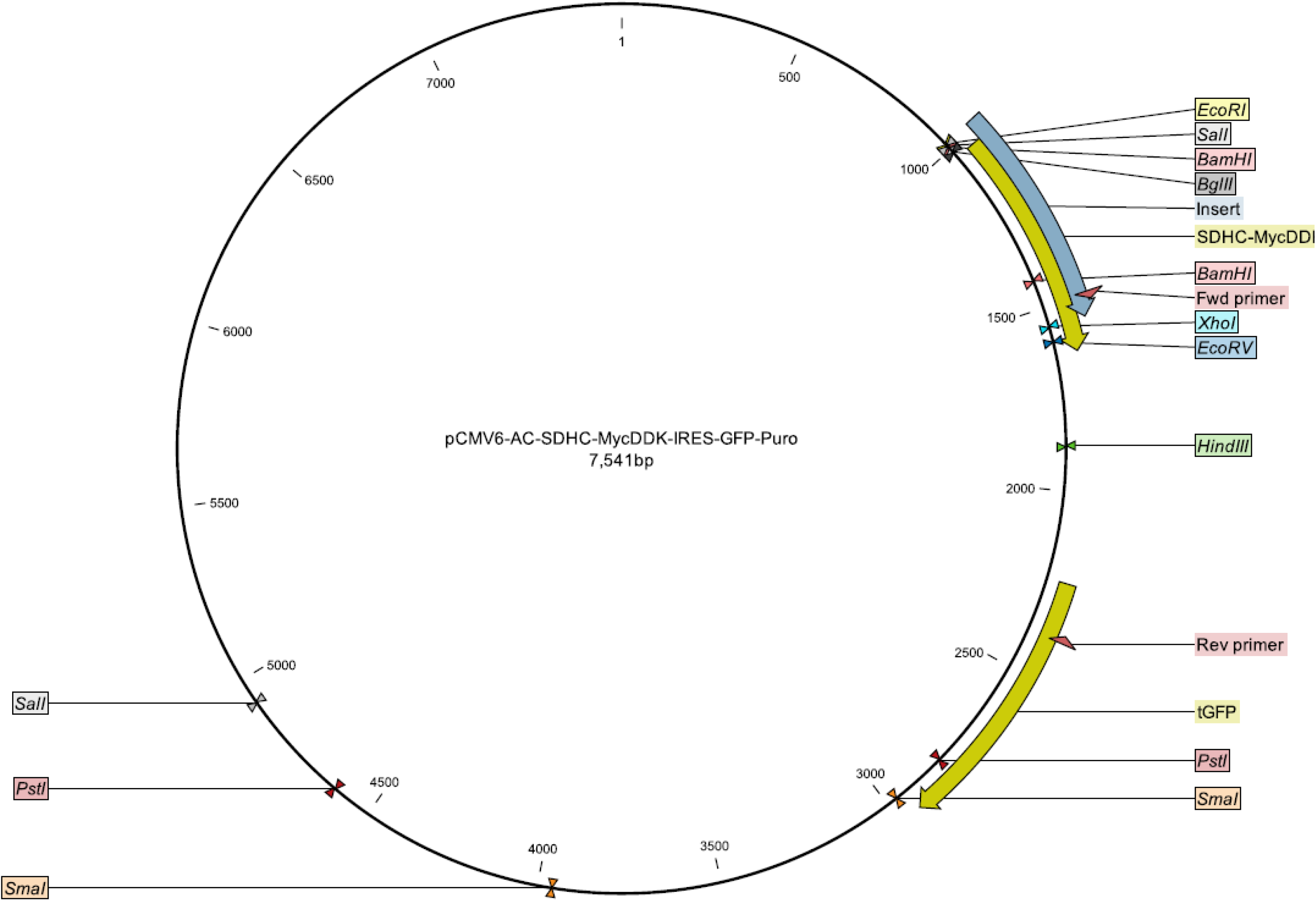
Plasmid map

**Figure S2:**
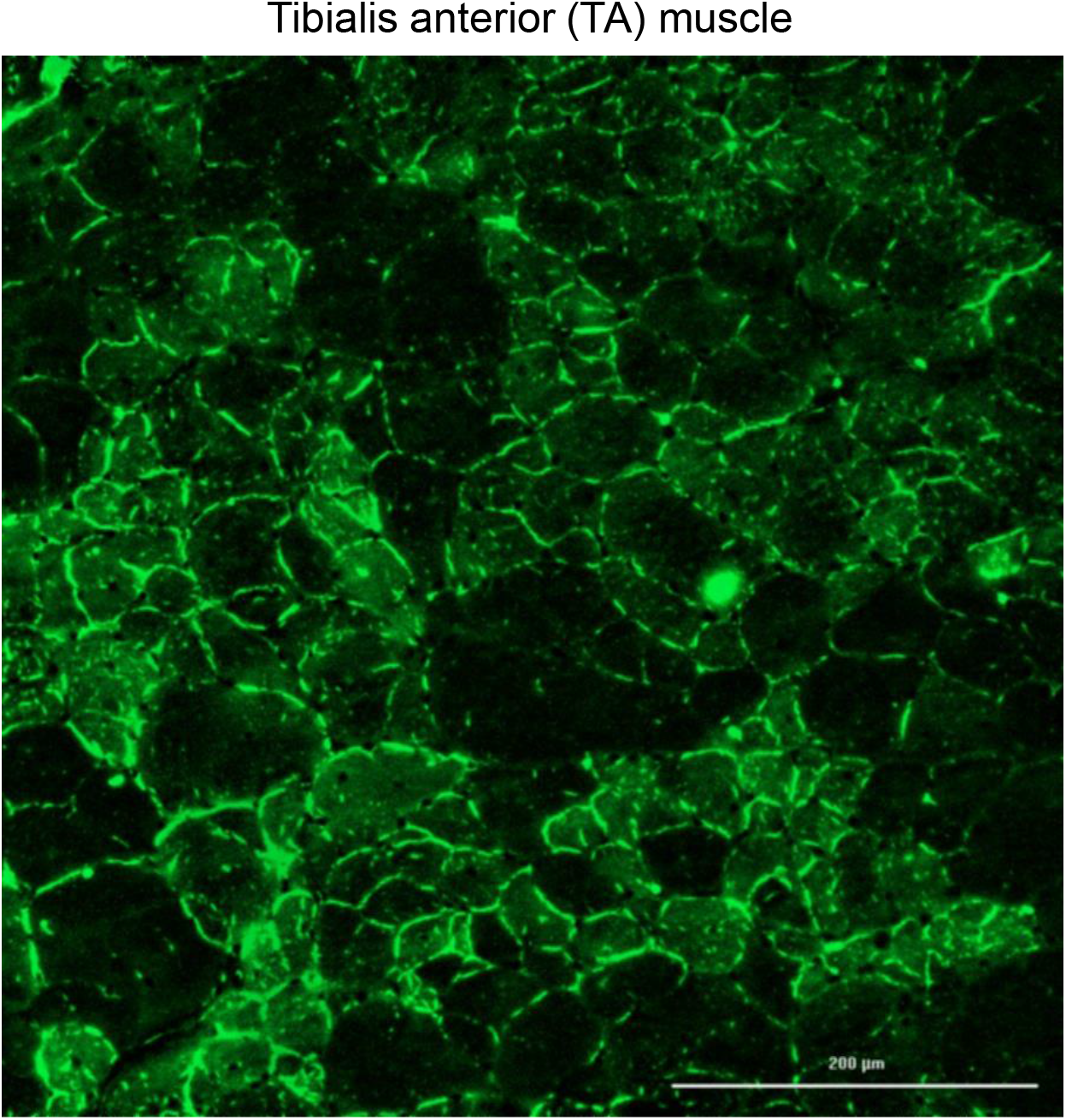
Fluorescence of TA electroporated muscle

**Figure S3:**
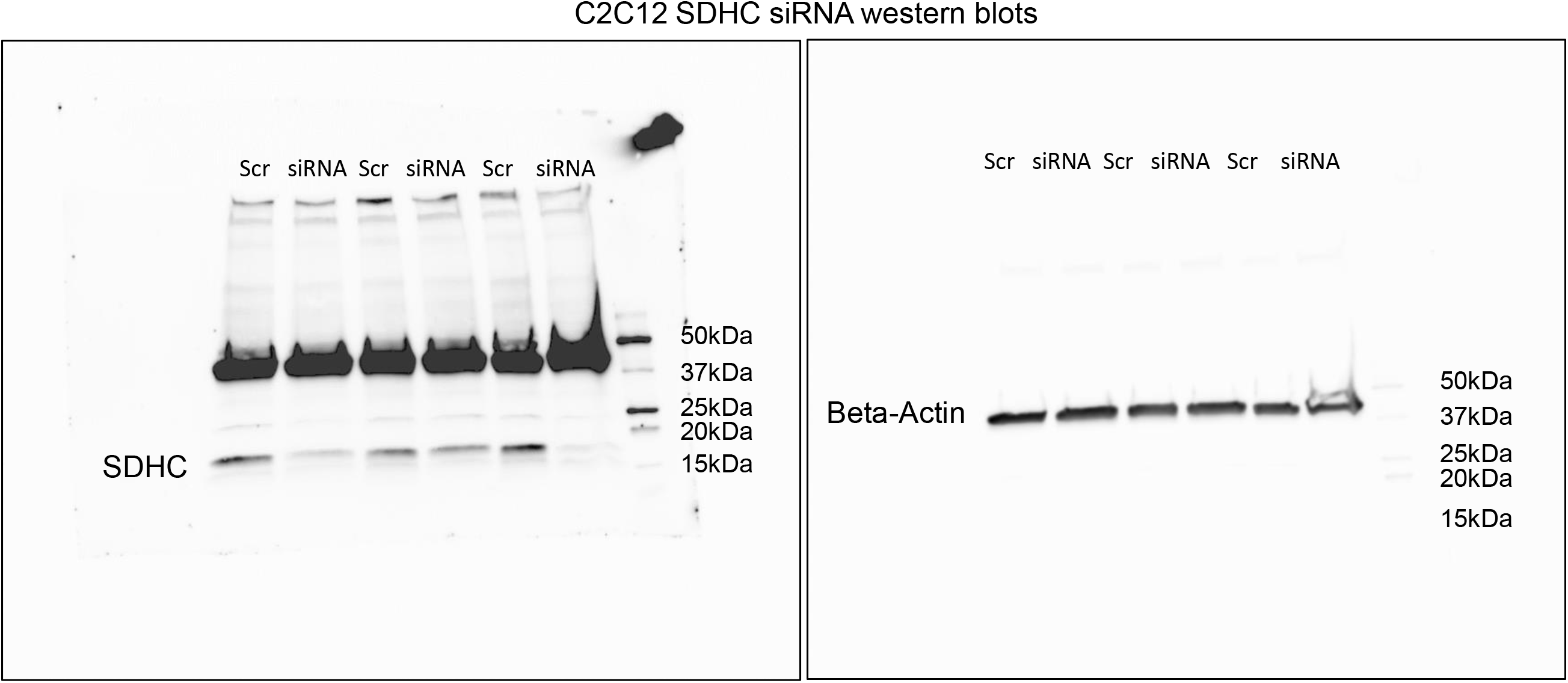
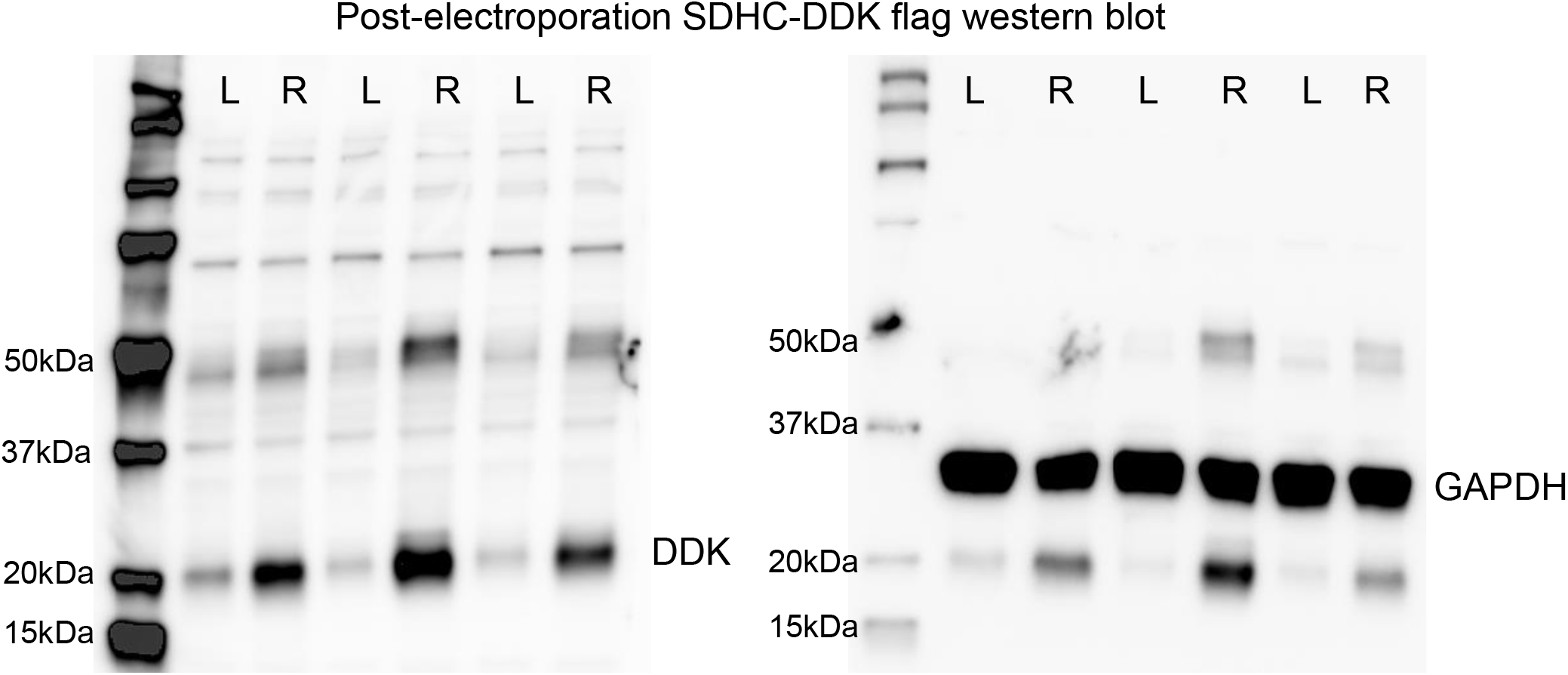
Original blots from flag and SDHC in C2C12 cells

**Figure S4:**
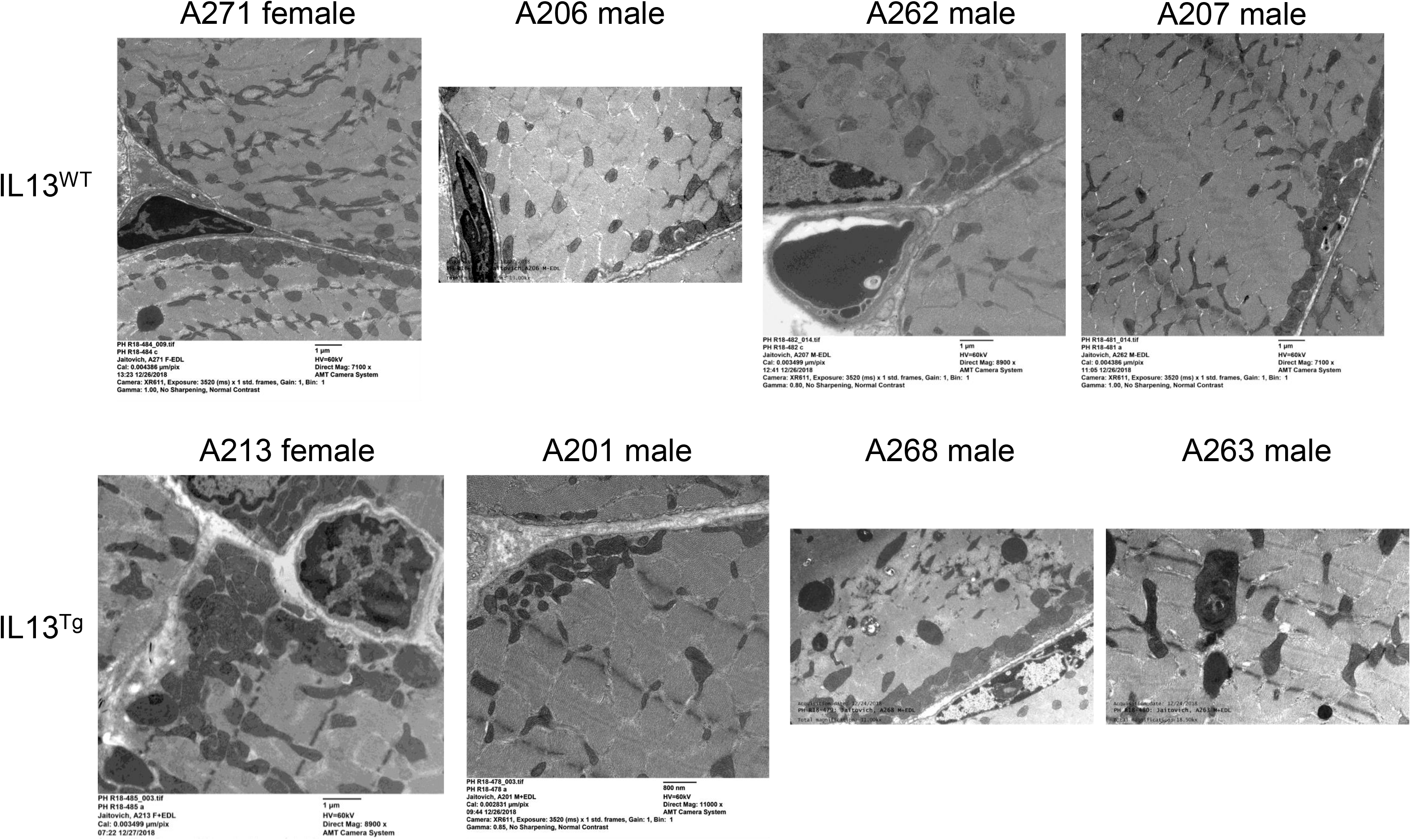
EM original pictures

**Table S1:** List of antibodies used in this project

| Target | Working concentration | Company (cat. no.) |
| --- | --- | --- |
| DDK-Flag | 1:1000 | Origene (TA50011) |
| SDHC | 1:1000 | Abcam (ab155999) |
| GAPDH | 1:3000 | Sigma (G9545) |
| Beta-Actin-HRP | 1:10000 | Thermo (MA5-15739-HRP) |
| Laminin | 1:100 | Sigma (L9393) |
| BF-F3 | 1:100 | DSHB (BA-D5, concentrate) |
| BA-D5 | 1:100 | DSHB (BA-D5, supernatant) |
| SC-71 | 1:100 | DSHB (SC-71, supernatant) |
| Anti-mouse IgG2b-DyLight 405 | 1:250 | Jackson IRL (115-475-207) |
| Anti-mouse IgG1-Alexa Fluor 488 | 1:250 | Jackson IRL (115-545-205) |
| Anti-rabbit Alexa Fluor 647 | 1:250 | Invitrogen (A-21245) |
| Anti-mouse Alexa Fluor 594 | 1:250 | Jackson IRL (115-585-020) |

**Table S2:**
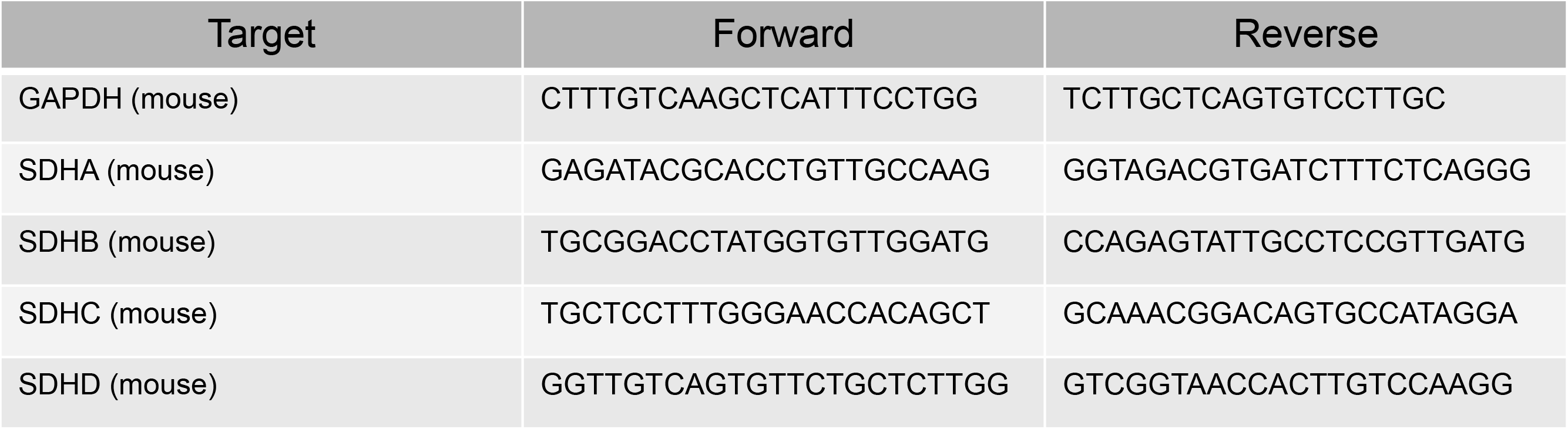
List of rtPCR primers used in this project

